# Retinal Ganglion Cell Axon Sorting at the Optic Chiasm Requires Dystroglycan

**DOI:** 10.1101/286005

**Authors:** Reena Clements, Kevin M. Wright

**Author notes:** Correspondence to Kevin M. Wright. Vollum Institute, 3181 SW Sam Jackson Park Rd L474 Portland, OR 97239.

## Abstract

In the developing visual system, retinal ganglion cell (RGC) axons project from the retina to several distal retinorecipient regions in the brain. Several molecules have been implicated in guiding RGC axons *in vivo*, but the role of extracellular matrix molecules in this process remains poorly understood. Dystroglycan is a laminin-binding transmembrane protein important for formation and maintenance of the extracellular matrix and basement membranes and has previously been implicated in axon guidance in the developing spinal cord. Using two genetic models of functional dystroglycan loss, we show that dystroglycan is necessary for correct sorting of contralateral and ipsilateral RGC axons at the optic chiasm. Missorted axons still target retinorecipient brain regions and persist in adult mice, even after axon pruning is complete. Our results highlight the importance of the extracellular matrix for axon sorting at an intermediate choice point in the developing visual circuit.

**Summary Statement:** Abnormal retinal ganglion cell axon sorting in the optic chiasm in the absence of functional dystroglycan results in profound defects in retinorecipient innervation.

## Introduction

The proper function of neural circuits is dependent on establishing precise connectivity during development. In the mammalian visual system, the axons of retinal ganglion cells (RGCs) leave the retina and cross the optic chiasm in the ventral forebrain *en route* to multiple retinorecipient areas of the brain. RGC axons terminate and form synapses in both image-forming brain regions, the lateral geniculate nucleus (LGN) and superior colliculus (SC), and non-image forming brain regions including the suprachiasmatic nucleus (SCN), olivary pretectal nucleus (OPN), nucleus of the optic tract (NOT) and medial terminal nucleus (MTN) (Zhang et al., 2017).

RGC axon guidance at the optic chiasm is dependent on several extracellular cues. Slit1 and Slit2 signaling through Robo receptors confines RGC axons to fasciculated tracts at the optic chiasm through surround repulsion (Plump et al., 2002). Vegf164/Nrp1, Sema3d/Nrp1 and Sema6D/NrCAM/PlxnA1 ligand/receptor complexes promote contralateral axon crossing at the chiasm (Dell et al., 2013; Erskine et al., 2011; Kuwajima et al., 2012; Sakai and Halloran, 2006). In the mouse, approximately 5% of neurons are excluded from the optic chiasm and form an ipsilateral projection in the optic tract (Petros et al., 2008). These ipsilateral axons are repelled by a combination of EphrinB2 expressed by optic chiasm glial cells, and Shh secreted transaxonally by contralateral RGC axons (Peng et al., 2018; Williams et al., 2003). Additional molecular cues govern axonal sorting, targeting of specific retinorecipient regions by subsets of RGC axons, and retinotopic mapping after crossing the chiasm (Feldheim et al., 1998; Hindges et al., 2002; Osterhout et al., 2011; Schmitt et al., 2006; Su et al., 2011; Suetterlin and Drescher, 2014; Sun et al., 2015).

In addition to canonical axon guidance molecules, components of the extracellular matrix (ECM) also play a critical role in axon guidance by generating both permissive and non-permissive substrates for axon growth. Throughout the nervous system, ECM proteins are highly enriched at basement membranes that form at the interface of neuroepithelial endfeet and the surrounding tissues. Basement membranes are highly dynamic structures that also regulate the extracellular localization of both attractive and repulsive secreted axon guidance cues (Dominici et al., 2017; Varadarajan et al., 2017; Wright et al., 2012; Xiao et al., 2011). The trajectory of RGC axons along the inner limiting membrane (ILM) keeps them in direct contact with the basement membrane as they exit the retina. However, whether the basement membrane regulates the trajectory of RGC axons through the optic chiasm has not been examined.

Dystroglycan is a highly glycosylated transmembrane protein that is critical for maintaining the structural integrity of basement membranes through its binding to several ECM proteins, including laminin and perlecan. Mutations in *dystroglycan* or one of the 18 genes required for its proper glycosylation result in compromised basement membrane integrity (Manya and Endo, 2017). This can lead to dystroglycanopathy, a form of congential muscular dystrophy that is frequently accompanied by neurodevelopmental defects that include type II lissencephaly and retinal dysplasia (Clements et al., 2017; Michele et al., 2002; Moore et al., 2002; Takeda et al., 2003; Taniguchi-Ikeda et al., 2016). Dystroglycan can also regulate axon guidance in the spinal cord by maintaining the basement membrane as a permissive growth substrate, and by restricting the localization of secreted Slit proteins to the floorplate. (Wright et al., 2012). Here, we identify a critical role for dystroglycan in regulating the sorting of RGC axons at the optic chiasm, resulting in inappropriate innervation patterns in retinorecipient regions in the brain.

## Results and Discussion

### Dystroglycan is required for axon sorting at the optic chiasm

In the mouse, after RGC axons exit the retina, approximately 95% cross at the optic chiasm (Petros et al., 2008). At e13, axons initially form a contralateral projection (Figure 1A, purple), followed by an ipsilateral projection (Figure 1A, green) forming between e14-e17 (Petros et al., 2008). Analysis of the ventral forebrain at e13 shows that dystroglycan protein is enriched in the basement membrane in direct proximity to where RGC axons cross to form the optic chiasm (Figure 1B). We therefore examined RGC axon trajectories in *ISPD*^*L79*/L79\**^ mutant mice, in which dystroglycan cannot bind to laminin or other ECM proteins due to defective glycosylation. These mice have intraretinal axon guidance defects due to degeneration of the retinal ILM and abnormal axonal projections in sensory afferents and commissural axons in the spinal cord (Clements et al., 2017; Wright et al., 2012). In control sections at e13, laminin staining is continuous along the ventral forebrain (Figure 1 C, E), with fasciculated bundles of RGC axons (Figure 1 D, E) growing in direct contact with the laminin. In contrast, laminin in the basement membrane is fragmented and discontinuous, and RGC axons appear defasciculated in *ISPD*^*79*/79\**^ mutants at e13 (Figure 1 C-E). Additional basement membrane proteins such as perlecan and collagen IV colocalize with laminin and are also discontinuous at the chiasm in mutants at e13 (Supplemental Figure 1A-D). Midline radial glia appear largely normal at e13 in *ISPD*^*L79*/L79\**^ mutants (Figure 1F), and there was no obvious structural defect at the chiasm.

**Figure 1:**
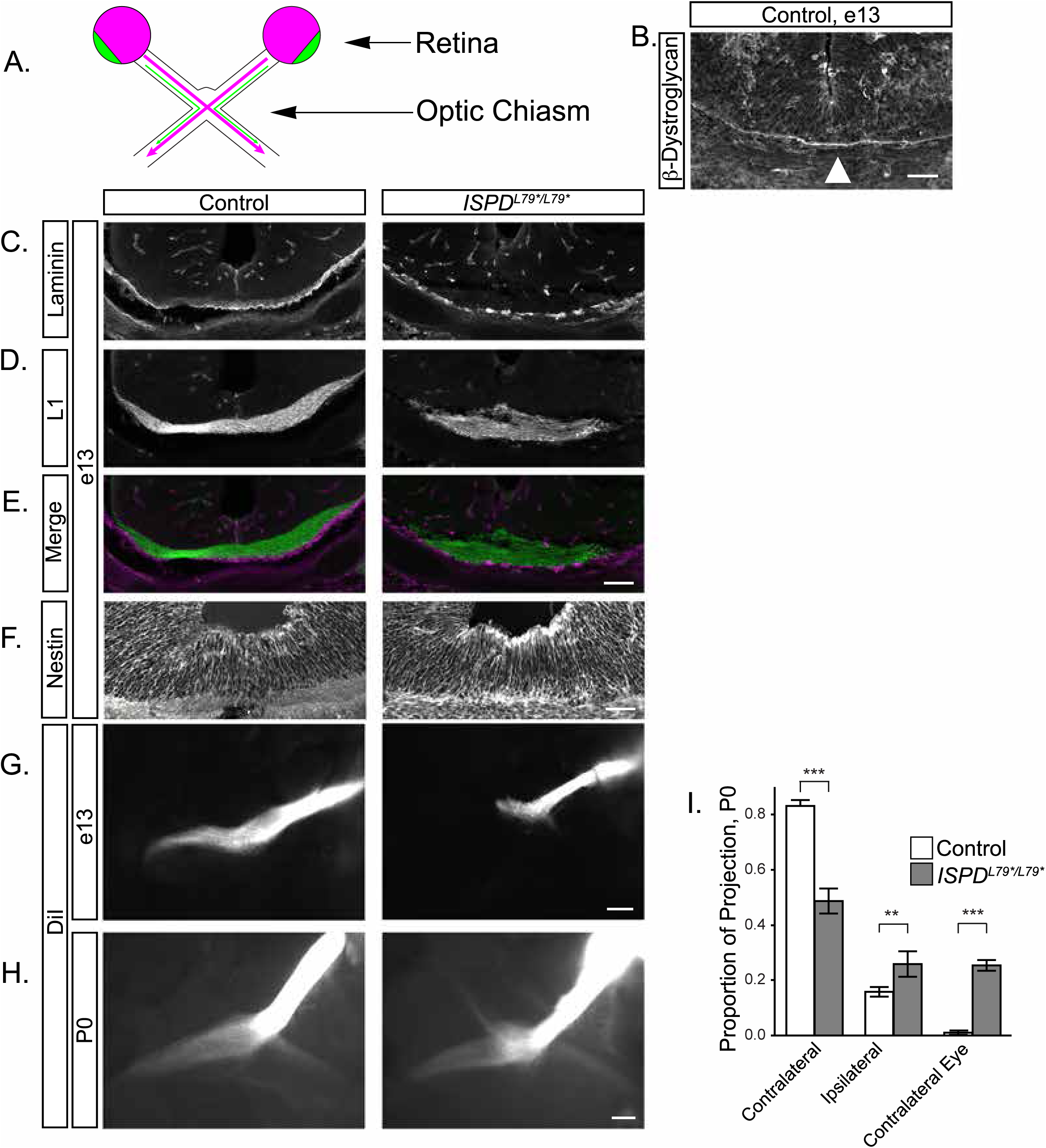
Glycosylated dystroglycan is required for axon guidance at the optic chiasm. (A) Schematic of RGC axon crossing at the optic chiasm. The contralateral projection is designated in purple, and the ipsilateral projection is designated in green. (B) At e13, β-Dystroglycan is enriched at the basement membrane at the chiasm (arrowhead). (C-E) Sections through the ventral forebrain at e13 reveal that RGC axons (L1, D, green, E) extend in direct proximity to the basement membrane (laminin, C, purple, E) in controls (left panels). The basement membrane is discontinuous in *ISPD*^*L79*/L79\**^ mutants (right panels), and axons are defasciculated and enter the ventral forebrain. (F) Midline glia at the optic chiasm labeled with nestin appear structurally intact in both controls and *ISPD*^*L79*/L79\**^ mutants. (G) At e13, DiI labeled RGC axons exclusively form a contralateral projection after crossing through the optic chiasm in controls (left). Axons in *ISPD*^*L79*/L79\**^ mutants fail to advance though the chiasm, and form an abnormal ipsilateral projection (right). (H) At P0, axons in controls make a dominant contralateral and minor ipsilateral projection (left). In *ISPD*^*L79*/L79\**^ mutants, axons project contralaterally, ipsilaterally, and to the contralateral eye (right). (I) Quantification of RGC axon trajectories indicate that mutant axons have a reduced contralateral projection and increased projection to both the ipsilateral optic tract and the contralateral eye (ANOVA, Tukey HSD *post hoc* test, p<0.01, n=12 control, 6 mutant chiasms). Scale bars: B, 50μm; C-E, 100μm; F, 50μm; G, H, 200μm.

In control mice, RGC axons at e13 labeled with intraretinal DiI extended through the optic chiasm to form a contralateral projection (Figure 1G). In contrast, in *ISPD*^*L79*/L79\**^ mutants, axons stall at the chiasm, fail to project into the contralateral optic tract, and form a premature projection into the ipsilateral optic tract (Figure 1G). By P0, many axons in *ISPD*^*L79*/L79\**^ mutants have progressed through the chiasm, but only 50% of RGC axons enter the contralateral optic tract, with the remainder of the axons inappropriately projecting into the ipsilateral optic tract and contralateral optic nerve (Figure 1 H, I, ANOVA, Tukey HSD *post hoc* test, p<0.01). Based on these results, we conclude that RGC axons fail to enter and sort appropriately at the optic chiasm in *ISPD*^*L79*/L79\**^ mutants.

To confirm that the basement membrane degeneration and axon sorting phenotypes we identified in *ISPD*^*L79*/L79\**^ mutants are due to loss of functional dystroglycan, we generated *DG*^*F/-*^; *Six3*^*Cre*^ mice to delete dystroglycan in both the retina and ventral forebrain (Figure 2A, B), (Furuta et al., 2000). We observed degeneration of the basement membrane in the ventral forebrain (Figure 2C, E), and axonal stalling and misrouting at the optic chiasm (Figure 2D, E, F) in *DG*^*F/-*^; *Six3*^*Cre*^ mice. These results verify that RGC axon guidance defects at the optic chiasm of *ISPD*^*L79*/L79\**^ mutants is indeed due to the loss of functional dystroglycan.

**Figure 2:**
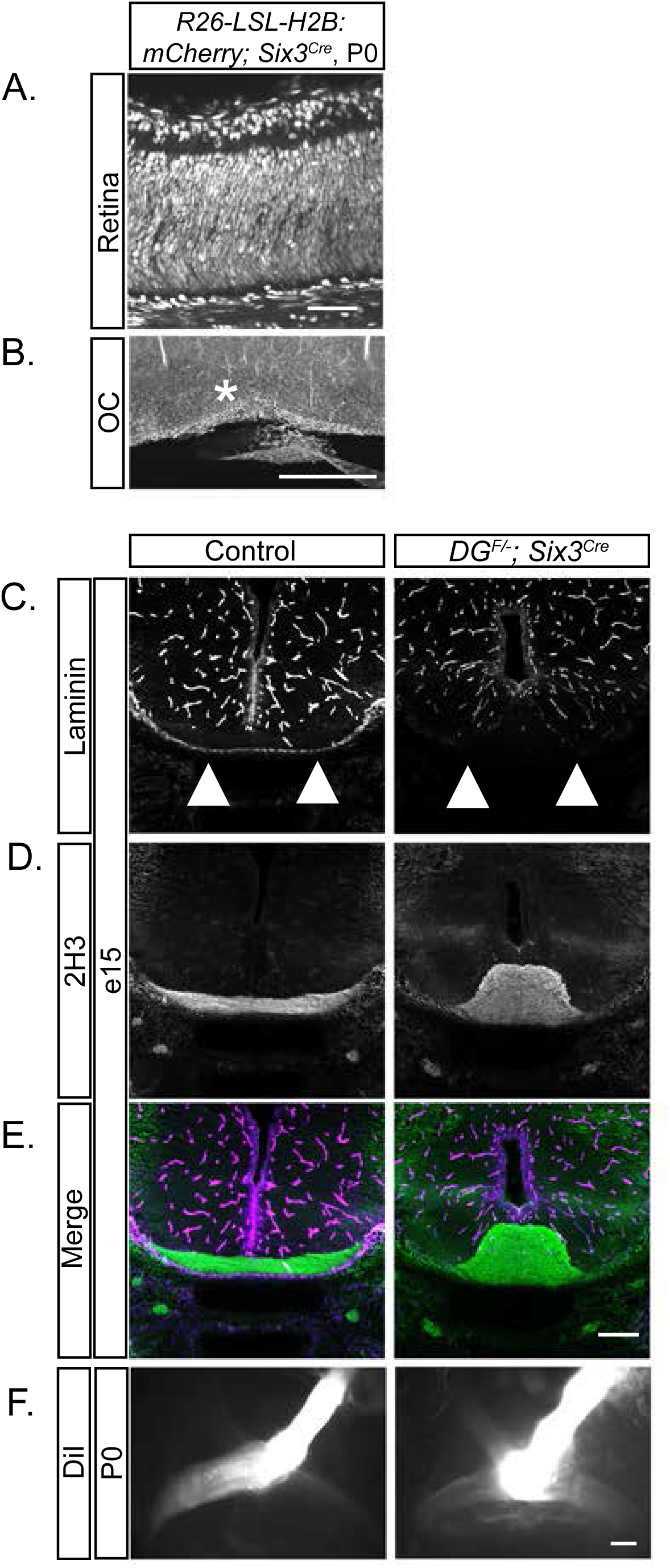
Conditional deletion of *dystroglycan* results in RGC axon guidance defects at the optic chiasm. (A-B) Sections from *Rosa26-lox-stop-lox-H2B:mCherry; Six3*^*Cre*^ mice at P0 show recombination throughout the retina (A) and in cells lining the optic chiasm. (C-E) Sections through the ventral forebrain at e15 reveal a loss of laminin staining at the chiasm (C, E purple) and axons (2H3) that inappropriately grow into the ventral forebrain (D, F green) in *DG*^*F/-*^; *Six3*^*Cre*^ mice. (F) RGC axons in *DG*^*F/-*^*;Six3*^*Cre*^ mutants stall at the optic chiasm and have reduced projections into the contralateral and ipsilateral optic tracts than in controls. Scale bars: A, 50 μm; B, 500μm; C-F, 200μm.

The disruption of basement membrane integrity in *ISPD*^*L79*/L79\**^ and *DG*^*F/-*^; *Six3*^*Cre*^ mutants suggests a potential mechanism for axonal misrouting. However, we also considered the possibility that dystroglycan could function cell-autonomously in RGCs. To test this, we generated *DG*^*F/-*^; *Isl1*^*Cre*^ conditional knockout mice, in which dystroglycan is deleted in all RGCs, but not the neuroepithelial cells that form the optic chiasm (Supplemental Figure 2A), (Pan et al., 2008). DiI labeling of RGCs did not reveal any differences in axon crossing between controls and *DG*^*F/-*^; *Isl1*^*Cre*^ mutants at P0 (Supplemental Figure 2B), suggesting that dystroglycan is not required in RGC axons to navigate the optic chiasm. We also examined whether ipsilaterally projecting RGCs were prematurely specified in the absence of dystroglycan. These cells are generated in the ventrotemporal retina at e14.5-18.5 and can be identified by the expression of the transcription factor Zic2 (Herrera et al., 2003). At e13, a timepoint when we first detected inappropriate projections into the ipsilateral optic tract in *ISPD*^*L79*/L79\**^ mutants, there were no Zic2+ RGCs (Supplemental Figure 3A, B). At e16, when the ipsilateral projection should be specified, the Zic2+ population appeared normal in mutants (Supplemental Figure 3C, D). Therefore, dystroglycan is not required for the proper specification of ipsilaterally projecting RGCs, and does not function within RGC axons for their proper sorting at the optic chiasm.

Based on these findings, we conclude that glycosylated dystroglycan is required specifically within the optic chiasm for the proper growth and sorting of RGC axons. While it is possible that the loss of dystroglycan affects the distribution of axon guidance cues at the chiasm, the defects we observe in *ISPD*^*L79*/L79\**^ and *DG*^*F/-*^; *Six3*^*Cre*^ mutants are far more severe than those observed following deletion of canonical axon guidance cues. Therefore, we favor a model in which dystroglycan is required to maintain basement membrane integrity, which functions as a permissive growth substrate for RGC axons at the optic chiasm.

### Axon mistargeting occurs within visual target regions due to loss of dystroglycan at the optic chiasm

After exiting the optic chiasm, RGC axons project to multiple retinorecipient regions of the brain. While *ISPD*^*L79*/L79\**^ mutants die perinatally, *DG*^*F/-*^; *Six3*^*Cre*^ mutants survive to adulthood, allowing us to examine how the loss of dystroglycan at the optic chiasm affects the ability of RGC axons to innervate their appropriate targets in the brain. Importantly, *Six3*^*Cre*^ does not drive a significant level of recombination in the LGN or SC (Supplemental Figure 4), allowing us to specifically examine how inappropriate sorting at the chiasm affects retinorecipient targeting.

To visualize axon tracts from each eye, we performed dual-color Cholera Toxin Subunit B (CTB) injections. At P2, the majority of axons in the optic tract posterior to the optic chiasm originate from the contralateral eye. However, axons from both the contralateral and ipsilateral retina were present in the optic tract in *DG*^*F/-*^; *Six3*^*Cre*^ mutants (Figure 3A). In the dLGN of control mice (Figure 3B), contralaterally projecting axons (purple) innervate the outer shell region, and ipsilateral axons (green) innervate the inner core. In stark contrast, *dystroglycan* mutant axons show extensive overlapping contralateral and ipsilateral innervation throughout the entire dLGN (Figure 3B). Although axons are still actively undergoing eye-specific segregation in the dLGN at P2, threshold-independent analysis indicates a statistically significant decreased variance of R in mutants, meaning that the projections have more overlap (Figure 3F, t-test, p=0.038) (Torborg and Feller, 2004).

**Figure 3:**
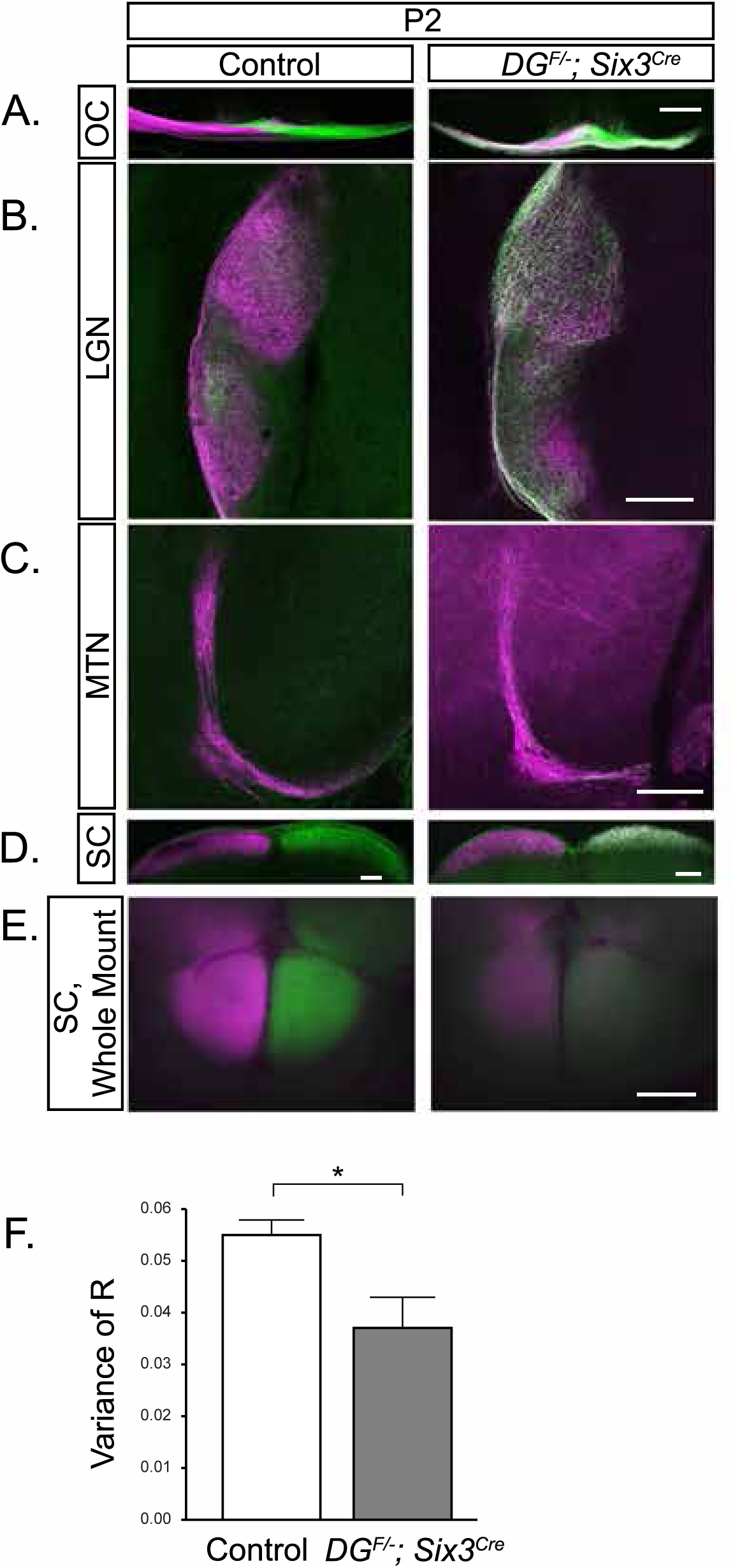
Retinal ganglion cell axons target appropriate visual regions in *DG^F/-^; Six3^Cre^* mice. (A) CTB labeled axons at the optic chiasm show eye-specific segregation in controls at P2 (left panel) and overlapping projections in *DG*^*F/-*^; *Six3*^*Cre*^ chiasms (right). (B) Contralateral axons (purple) innervate the dorsal and lateral LGN while ipsilateral axons (green) begin to grow into the dorsal and lateral LGN cores in controls (left) at P2. Axons originating from contralateral and ipsilateral eye overlap across the entirety of the LGN in *DG*^*F/-*^; *Six3*^*Cre*^ mutants. (C) In the MTN of the accessory optic system, controls receive monocular innervation from the contralateral eye (left), while the MTN in *DG*^*F/-*^; *Six3*^*Cre*^ mutants contains ipsilaterally originating axons. (D, E) In sections (D) and whole mount views of the superior colliculus (E), controls (left) are predominantly innervated by the contralateral eye. In *DG*^*F/-*^; *Six3*^*Cre*^ mice (right), there is extensive innervation from both contralateral and ipsilateral eyes across the entire span of the colliculus. (F) Threshold-independent analysis of LGN projections (B) shows a significantly greater axonal overlap in the dLGN in mutants than controls (t-test, p=0.038, n=3 control, 3 mutant dLGNs). Scale bar 200μm.

In control mice, the SC receives innervation primarily from the contralateral retina (Figure 3D, E). In *DG*^*F/-*^; *Six3*^*Cre*^ mutants, axons correctly target the SC, but exhibit overlapping contralateral and ipsilateral projections spanning the entire SC (Figure 3D, E). The reduced intensity of CTB labeled RGC projections in the colliculus (Figure 3E) likely reflects increased apoptotic death of RGCs in *DG*^*F/-*^; *Six3*^*Cre*^ mutants, which may arise from axons that fail to exit the optic chiasm (Clements et al., 2017). RGC axons in *DG*^*F/-*^; *Six3*^*Cre*^ mutants also innervate the MTN, a component of the non-image forming visual system (Figure 3C). Thus, RGC axons specifically innervate retinorecipient brain regions of the brain and do not project into inappropriate brain regions in *DG*^*F/-*^; *Six3*^*Cre*^ mutants. However, the innervation of each retinorecipient region is abnormally binocular.

In the LGN and SC, RGC axons undergo refinement prior to eye opening to complete their specific laminar and retinotopic organization (Godement et al., 1984; Jaubert-Miazza et al., 2005). This refinement is mediated by spontaneous activity in the form of retinal waves, Eph/Ephrin signaling, and caspase-dependent pruning (Butts et al., 2007; Feldheim et al., 1998; Hindges et al., 2002; McLaughlin et al., 2003; Simon et al., 2012). We have previously shown that spontaneous retinal waves are preserved in *dystroglycan* null mice (Clements et al., 2017). To determine whether the inappropriate ipsilateral axons observed at P2 were eliminated by axon pruning, we performed CTB injections prior to eye opening at P10, and at P28, when eye-specific axon segregation is complete (Furman and Crair, 2012). Surprisingly, while pruning does occur after P2, ipsilateral axons that have inappropriately invaded contralateral retinorecipient regions persist into adulthood. At both P10 (Figure 4A) and P28 in *DG*^*F/-*^; *Six3*^*Cre*^ mutants (Figure 4E), we observe axons from both the ipsilateral and contralateral eyes in the optic tract within the optic chiasm. In the LGN of *DG*^*F/-*^; *Six3*^*Cre*^ mutants, (Figure 4B, F), axons fail to prune in a manner that correctly forms a contralateral shell and ipsilateral core. Rather, ipsilateral axons are present in patchy, random distributions throughout the entire LGN. To determine if the appropriate proportion of contralateral and ipsilateral axons is maintained in the dLGN despite their inappropriate innervation pattern, we measured the area occupied by contralateral, ipsilateral, and overlapping axons. We observed a significant increase in the proportion of ipsilateral axons in *DG*^*F/-*^; *Six3*^*Cre*^ mutants at P10 (Figure 4I, p=0.04, t-test), whereas the proportion of contralateral and overlapping projections do not show any difference (Figure 4I, p>0.05, t-test). Axons in the dLGN continued pruning between P10 and P28, as the proportion of ipsilateral projections was not statistically different between *DG*^*F/-*^; *Six3*^*Cre*^ mutants and controls at P28 (Figure 4I, p>0.05, t-test). Using a threshold-independent method to measure segregation between contralateral and ipsilateral projections, we found that despite ipsilateral projections innervating inappropriate regions of the dLGN shell, contralateral and ipsilateral projections still exhibit normal eye-specific segregation (Figure 4J, p>0.05, t-test). The ability of misprojecting axons to undergo a normal process of segregation is likely due to the presence of retinal waves in *DG*^*F/-*^; *Six3*^*Cre*^ mutants.

**Figure 4:**
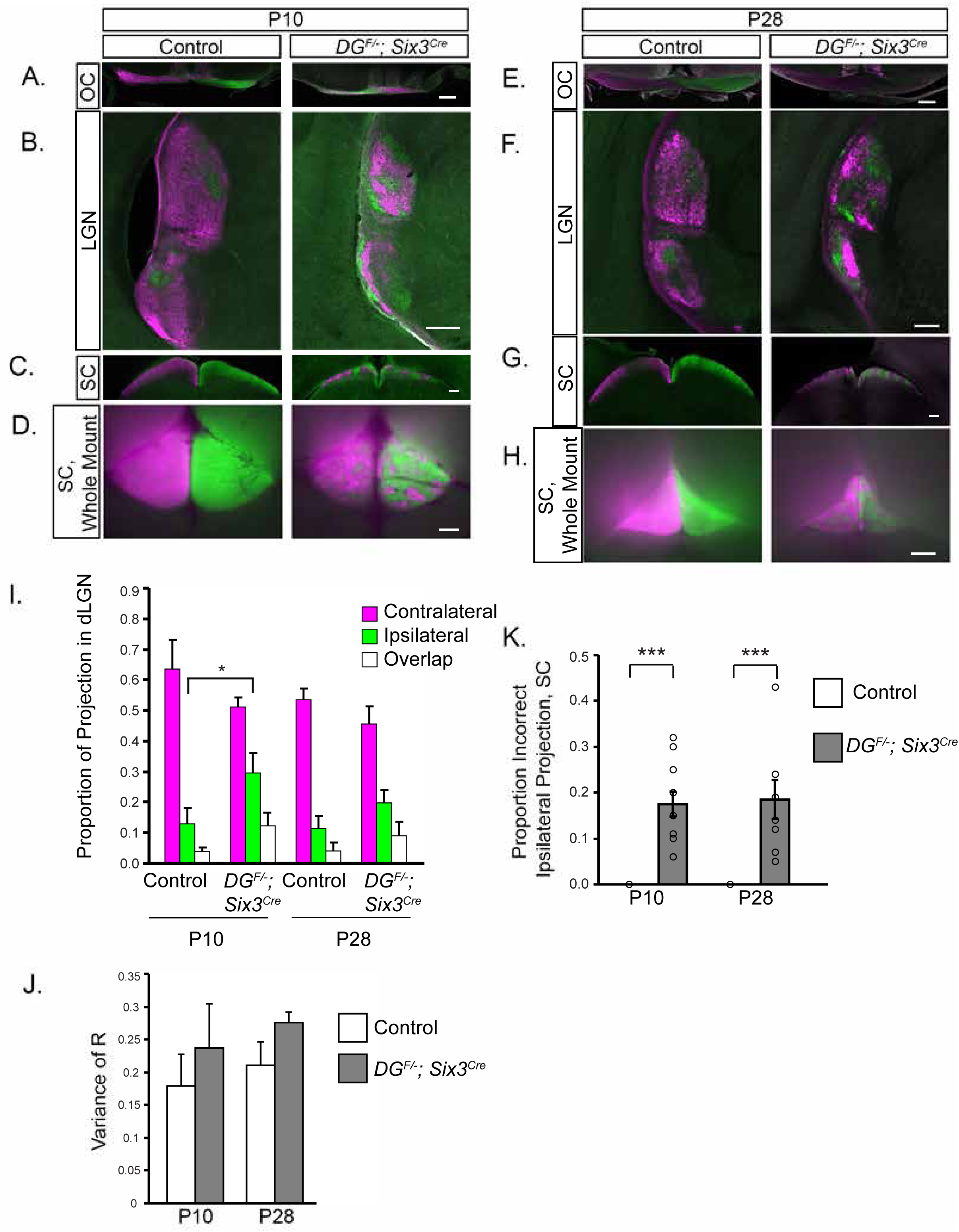
Abnormal RGC innervation of retinorecipient regions persists into adulthood in *DG^F/-^; Six3^Cre^* mice. (A) CTB labeling of RGC axons reveals extensive overlap between contralateral and ipsilateral axons at the optic chiasm in *DG*^*F/-*^; *Six3*^*Cre*^ mice at P10. (B) CTB labeling shows that while RGC pruning occurs in *DG*^*F/-*^; *Six3*^*Cre*^ P10 dorsal and lateral LGN, axons do not prune into the appropriate contralateral shell/ipsilateral core arrangement. (C, D) SC sections (C) and whole mounts (D) in P10 *DG*^*F/-*^; *Six3*^*Cre*^ mice reveal extensive ipsilateral innervation that prunes into discrete patches. (E-H). Analysis of CTB injected axons in the optic chiasm (E), LGN (F), and SC (G, H) at P28 in *DG*^*F/-*^; *Six3*^*Cre*^ show that inappropriate ipsilateral projections remain in each of these areas after eye opening. (I) Threshold-dependent measurement of the total area of contralateral, ipsilateral, and overlapping projections in the dLGN show that the amount of ipsilateral innervation is significantly greater in *DG*^*F/-*^; *Six3*^*Cre*^ mice at P10 (p=0.04, t-test, n=4 controls, 4 mutants at P10, n=4 controls, 4 mutants at P28), but not statistically different from controls at P28. (J) Threshold-independent analysis does not show a difference in contralateral/ipsilateral segregation between controls and mutants at P10 or P28 (t test, p>0.05, n=4 controls, 4 mutants at P10, n=4 controls, 4 mutants at P28). (K) In *DG*^*F/-*^; *Six3*^*Cre*^ mice, approximately 20% of the axons in the SC at P10 and P28 inappropriately arise from the ipsilateral eye (ANOVA, Tukey HSD *post hoc* test, p<0.0001, n=7 control and 6 mutant SCs at P10, n=4 control and 4 mutant SCs at P28). Scale bars: A-C, E-G, 200μm; D, H, 500μm.

In *DG*^*F/-*^; *Six3*^*Cre*^ mutants (Figure 4C, D, G, H), clusters of ipsilateral axons abnormally invade the SC, evident in both sections and whole mounts. Analysis of whole mount SC at P10 and P28 reveals that approximately 20% of the innervation in the SC of *DG*^*F/-*^; *Six3*^*Cre*^ mutants originates from the ipsilateral eye, which persists into adulthood (ANOVA, Tukey HSD *post hoc* test, p<0.0001, Figure 4K). Therefore, the inappropriate ipsilateral projections that are distributed throughout the superior colliculus at P2 undergo pruning, resulting in their segregation from contralateral projections.

In addition to the dLGN and SC, abnormal innervation was found in non-image forming regions. For example, the MTN, which is normally monocularly innervated, receives binocular innervation in *dystroglycan* mutant mice (Supplemental Figure 5C, F). The SCN, which normally receives input from both eyes, continued to receive bilateral innervation (Supplemental Figure 5A, D). The optic tract, which contains axons from both eyes in controls and mutants, appeared markedly thinner in *DG*^*F/-*^; *Six3*^*Cre*^ mutants at both P10 and P28 (Supplemental Figure 5B, E), presumably due to increased RGC apoptosis previously described in these mice (Clements et al., 2017).

We can draw three conclusions from the CTB labeling data. First, in contrast to other models with optic chiasm guidance defects, such as Slit/Robo mutants, RGC axons in *dystroglycan* mutants only innervate retinorecipient regions and do not misproject into other areas of the brain. Second, the increased axonal overlap that we initially observe at P2 in both the dLGN and SC of *DG*^*F/-*^; *Six3*^*Cre*^ mutants undergoes pruning, resulting in complete segregation of axons from contralateral and ipsilateral eyes at later stages. This pruning presumably relies on spontaneous activity in the retina, as much of it occurs by P10, prior to eye opening. Third, despite the segregation of contralateral and ipsilateral axons, ipsilateral axons innervating inappropriate regions of the dLGN shell and SC fail to undergo elimination and persist into adulthood. These findings raise interesting questions of how an abnormal persistence of ipsilateral innervation in the dLGN and SC might affect downstream components of the visual circuit in *DG*^*F/-*^; *Six3*^*Cre*^ mutants. First, how does the presence of ipsilateral RGC axons in the shell region of the dLGN affect the formation of ocular dominance columns in V1 of the cortex? Second, does the abnormal pattern of ipsilateral RGCs affect other axons innervating the same structures? Two recent studies show that retinogeniculate and corticothalamic axons interact and may rely on one another for correct innervation of the LGN (Seabrook et al., 2013; Shanks et al., 2016). Finally, what is the impact of persistent ipsilateral innervation in inappropriate regions of the dLGN and SC on visual perception? Unfortunately, since dystroglycan is required for transmission at photoreceptor synapses, we are unable to test these questions in *DG*^*F/-*^; *Six3*^*Cre*^ mutants (Sato et al., 2008).

In summary, we show that dystroglycan plays a critical role in regulating the guidance and sorting of RGC axons at the optic chiasm by maintaining the integrity of the basement membrane. The loss of basement membrane integrity in the ventral forebrain of *ISPD*^*L79*/L79\**^ and *DG*^*F/-*^; *Six3*^*Cre*^ mutants coincident with the arrival of RGC axons at the chiasm suggests that this is the underlying cause of axonal misrouting. In *DG*^*F/-*^; *Six3*^*Cre*^ mutants, mis-sorted ipsilateral RGC axons continue to innervate inappropriate regions of retinorecipient regions of the brain into adulthood. Coupled with the requirement for dystroglycan at photoreceptor synapses and its role in regulating neuronal migration and dendritic lamination in the retina, our result provide insight into potential mechanisms that underlie visual impairments in patients with dystroglycanopathy.

## Materials and Methods

### Animals

Animal procedures were approved by the OHSU Institutional Animal Care and Use Committee and conformed to the National Institutes of Health *Guide for the care and use of laboratory animals*. Embryonic day 0 (e0) was the day of vaginal plug observation and postnatal day 0 (P0) was the day of birth. Animals were euthanized with CO_2_. Controls were *ISPD*^*+/L79\**^, and *DG*^*F/+*^; *Six3*^*Cre*^ or *DG*^*F/+*^ age matched littermates. *ISPD*^*L79*/L79\**^ (Wright et al., 2012) and *DG*^*F/F*^ (Moore et al., 2002) mice and genotyping have been described previously. Generic cre primers were used for *Six3*^*Cre*^ (Furuta et al., 2000) and *Isl1*^*Cre*^ (Yang et al., 2006) genotyping.

### Statistical Analysis

All analysis was performed on n≥3 mice from at least two litters. Statistics were conducted using JMP Pro 13.0 software (SAS Institute). A Student’s t-test was used to compare means of two groups, and an ANOVA with Tukey post-hoc test was used to compare means of two or more groups. A significance threshold of 0.05 was used for all statistical tests. * indicates p<0.05; ** indicates p<0.01; *** indicates p<0.001.

### Labeling and Analysis of the Optic Chiasm

DiI injections were performed at e13 and P0. Animals were decapitated and heads were fixed in 4% PFA at 4°C overnight. The left eye was enucleated and a crystal of DiI approximately the size of the optic disc was placed inside the optic disc. Low melting point 2.5% agarose was placed into the eyecup to hold the crystal in place. Heads were incubated at 37°C for three days at e13, one week at P0. Projections were visualized by removing the lower jaw, tongue, and palate to expose the ventral forebrain. Images were acquired on a Zeiss AxioZoom.V16.

Analysis of DiI projections at P0 (Figure 1) was conducted by tracing the area of the projections in FIJI and dividing by the total area of all projections for each mouse.

### Labeling and Analysis of Retinorecipient Brain Regions

Cholera Toxin Subunit B (CTB) injections to label anterograde axonal projections were performed at P0 or P1 for P2 analysis, at P0-P4 for P10 analysis, and at P26 for P28 analysis. Mice between P0-P4 were anesthetized on ice, and mice aged P26 were anesthetized with isofluorane. Glass capillary needles were backfilled with CTB conjugated to Alexa-488 or Alexa-555. The right eye was injected with CTB-488 and the left eye was injected with CTB-555. Injections were performed using a picospritzer injecting for 15 milliseconds at 30 PSI. At P2 or P10, brains were removed and drop fixed in 4% PFA at 4°C overnight. At P28, animals were perfused, and brains were post-fixed in 4% PFA at 4°C overnight. Whole mount images of the superior colliculus were obtained prior to fixation on a Zeiss AxioZoom.V16. After fixation, brains were washed in PBS for 30 minutes, embedded in 2.5% low melting point agarose, and sectioned on a vibratome at 150 μm. Sections were incubated in DAPI overnight and mounted using Fluoromount medium. Brain sections were imaged on a Zeiss Axio Imager M2 upright microscope equipped with an ApoTome.2. Analysis of superior colliculus misprojections (Figure 4K) was conducted on whole mount colliculus images by tracing the regions with inappropriate ipsilateral projections, adding to determine total area, and dividing by the area of the superior colliculus.

A threshold-dependent analysis method was used to determine the proportion of contralateral and ipsilateral projections to the dLGN based on previously described methods (Jaubert-Miazza et al., 2005). Briefly, the LGN was thresholded in FIJI to determine the pixels corresponding to either contralateral signal or ipsilateral signal. Threshold values were kept constant between corresponding contralateral and ipsilateral images. The total area of pixels in each image, and the overlap between the two areas, was measured.

A threshold-independent method was utilized to confirm results and analyze segregation between contralateral and ipsilateral projections (Torborg and Feller, 2004). We applied a rolling ball filter of 200 pixels to the LGN to subtract background. The logarithm of the intensity ratio (R) was calculated for each pixel to compare the intensity of ipsilateral (F_I_) and contralateral (F_C_) fluorescence, where R = log_10_(F_I_/F_C_). This generates a distribution of R-values for each LGN image. We then computed the variance for each R-distribution. Smaller variances reflect greater overlap, and less segregated axonal input, whereas larger variances reflect decreased overlap and more segregated axonal input.

Brains containing cre-dependent fluorescent markers used to define regions of cre recombination in retinorecipient brain regions were processed the same way as brains injected with CTB.

### Immunohistochemistry

Embryos used for immunohistochemistry were decapitated and fixed in 4% PFA at 4°C overnight. Tissue was washed in PBS for 30 minutes and equilibrated in 15% sucrose at 4°C overnight. Tissue was flash frozen in embedding media and sectioned at 20 μm on a cryostat. Sections were blocked with PBS containing 2% Normal Donkey Serum and 0.2% Triton for 30 minutes, and incubated in antibody diluted in this solution at 4°C overnight. Tissue was washed for 30 minutes and incubated in secondary antibody diluted in a PBS solution containing 2% Normal Donkey Serum for 2-4 hours. Tissue was incubated in PBS containing DAPI for 10 minutes, washed in PBS for 20 minutes, and mounted using Fluoromount medium. Sections were imaged on a Zeiss Axio Imager M2 upright microscope equipped with an ApoTome.2. Preparation of P0 retinas (Supplemental Figure 2) was performed as previously described (Clements et al., 2017). Antibody information is contained in Supplemental Table 1.

## Acknowledgements

We would like to thank Chinfei Chen and Genelle Rankin for providing software and assistance performing threshold-independent analysis of LGN innervation and members of the Wright laboratory for insightful discussion and advice throughout the course of this study. This study was funded by the National Institutes of Health Grant R01-NS091027 (K.M.W.), The Whitehall Foundation (K.M.W.), The Medical Research Foundation of Oregon (K.M.W.), National Science Foundation Graduate Research Fellowship Program (R.C.), and LaCroute Neurobiology of Disease Fellowship (R.C.).

## Supplemental Table 1: Antibodies

**Table 1:**
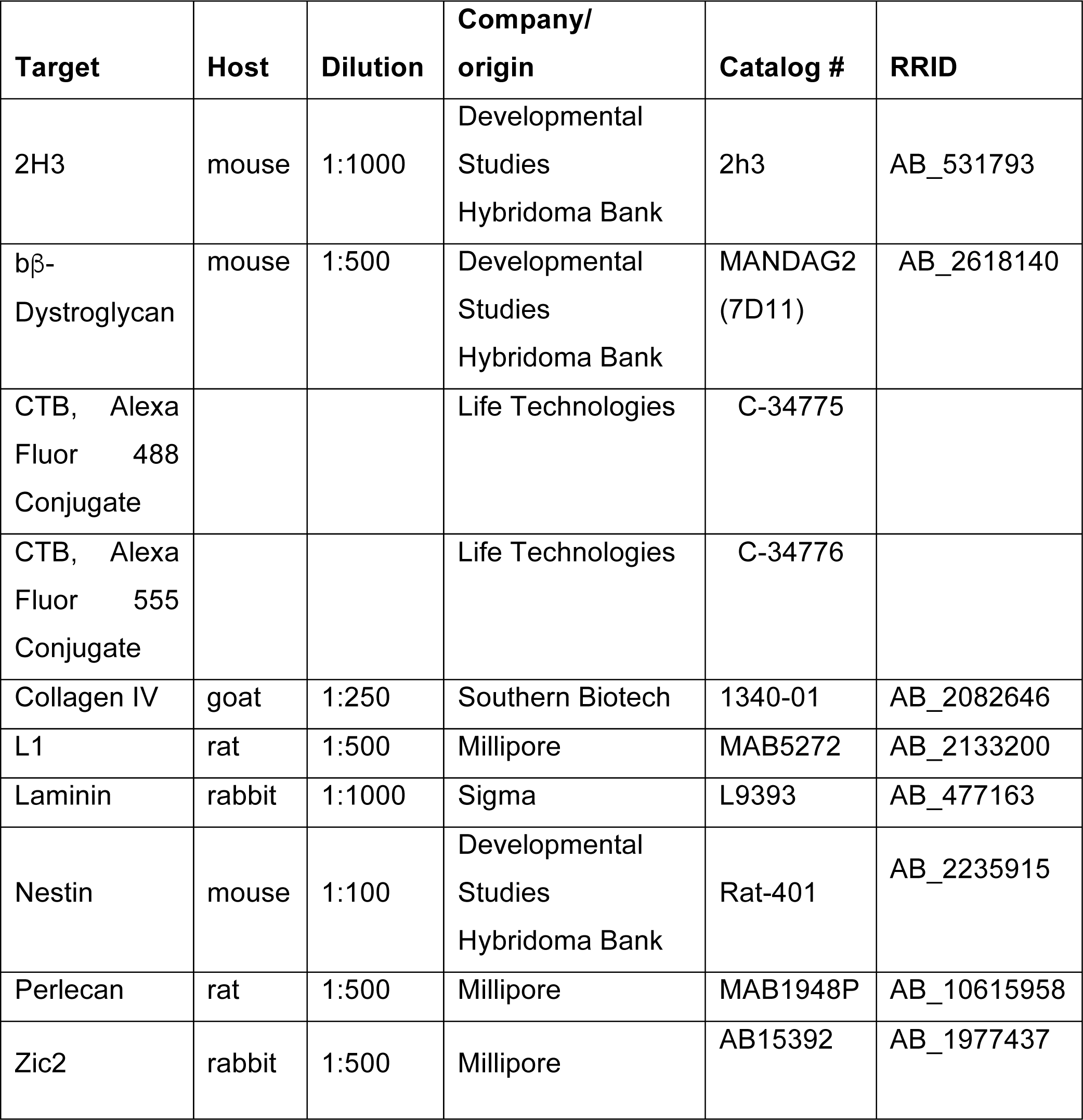
List of Antibodies. Antibodies and associated information (host species, dilution, manufacturer, catalog number, and RRID) are presented here.

